# Spatial transcriptomics reveals distinct inflammatory and adaptive immune properties of rheumatoid arthritis synovial fibroblasts

**DOI:** 10.1101/2025.11.08.687060

**Authors:** Camilla R.L. Machado, Mina Yao, David L. Boyle, Robert J. Benschop, James T. Parker, Wei Wang, Gary S. Firestein

## Abstract

**Objective:** Rheumatoid arthritis (RA) synovium displays cellular heterogeneity, with gene expression driving pathogenesis. Prior transcriptomic studies relied on disaggregated tissue, which causes cell loss and induction. We applied spatial transcriptomics to investigate synovial lining and sublining fibroblasts and macrophages from RA and osteoarthritis (OA) patients.

**Methods:** Fresh frozen synovial tissues from 7 RA and 8 OA patients were analyzed using the NanoString GeoMX DSP Whole Transcriptome Assay. Lining and sublining regions were segmented into fibroblasts and macrophages. Principal component analysis separated samples by cell type and disease. Differentially expressed genes (DEG) were analyzed by linear mixed models (p-value<0.05 and |log2 fold change|>0.5) and Reactome pathway (FDR<0.02) analysis.

**Results:** DEG analysis of RA compared to OA revealed distinct gene signatures across regions and cell types. RA lining fibroblasts exhibited a strong pro-inflammatory and matrix-destructive signature, while RA sublining fibroblasts showed an unexpected role in antigen presentation and adaptive immunity. RA lining macrophages exhibited enrichment for interleukin signaling, extracellular matrix organization, and translation-related pathways. In contrast, sublining macrophages showed minimal transcriptional differences between RA and OA, suggesting limited pathogenic involvement. Comparison between lining and sublining within RA showed that lining fibroblasts display a higher activated phenotype than sublining cells.

**Conclusion:** Spatial transcriptomic analysis uncovers distinct region- and cell-type-specific transcriptional profiles in RA synovium. Lining fibroblasts are highly activated and destructive, and sublining fibroblasts contribute surprisingly to adaptive immunity. This data provides clues to region-cell-type-specific functions that could be exploited to identify novel therapeutic targets.

## Introduction

Rheumatoid arthritis (RA) is a chronic immune-mediated disease characterized by synovial inflammation and progressive bone and cartilage destruction (1–3). The rheumatoid synovium is marked by heterogeneous infiltration of adaptive and innate immune cells that express a variety of inflammatory mediators that drive the process (4, 5). Although significant advances in understanding disease pathogenesis and the introduction of targeted therapies have revolutionized RA therapy, there remains a major unmet medical need because synovitis is not fully controlled in many patients (2, 4, 6). Thus, the cellular and regional mechanisms that maintain inflammation are still not fully understood.

The RA synovium is a complex tissue that is comprised of an intimal lining layer and a deeper sublining, each with distinct cell populations and functions. The lining region is composed of specific fibroblasts and macrophages, while the sublining is composed of other fibroblasts, a variety of immune cells, and blood vessels (6, 7). Rheumatoid lining fibroblasts have an aggressive phenotype, which promotes synovitis, pannus growth, and cartilage/bone destruction (3). These cells interact with neighboring lining macrophages, which produce their own panel of mediators (7, 8). Understanding transcriptional programs of synovial fibroblasts and macrophages in the lining and sublining could enhance improve disease subclassification and patient stratification for therapy.

Studies with unbiased cell-specific transcriptomes in RA synovium, especially compared with OA, have previously relied primarily on disaggregated tissues to understand the RA pathogenic processes (9–11). They revealed subpopulations of immune and mesenchymal cells associated with RA by using high-dimensional integration of single-cell RNA sequencing (RNAseq) with mass cytometry. Another study using disaggregated synovium tissue, laser capture microdissection (LCM), and single-cell RNAseq, observed transcriptional differences between RA and OA synovial tissue in lining and sublining (12). However, there are some limitations of using the tissue disaggregation, such as loss of tissue architecture, cell loss and gene induction (9). Moreover, there is a need for complementary methods for studies of gene expression using techniques that preserve the synovial cellular relationships, such as spatial transcriptomics (13). In this study, we interrogated RA and OA synovial tissue using spatial transcriptomics to investigate regional and cell-type differences, focusing on lining and sublining regions segmented into fibroblasts and macrophages. The results demonstrate unique topographical and cell lineage-specific gene expression patterns that likely contribute to the pathogenesis of RA.

## Methods

### Synovial tissue

Synovial tissue was obtained from patients with RA or OA undergoing arthroplasty with informed consent under Human Research Protection Program approval (protocol 140175). Patients with RA met the American College of Rheumatology 2010 criteria, and for OA, the diagnosis conformed to the 1991 criteria. Each specimen was dissected, embedded in ideal cutting temperature (OCT) medium, snap-frozen, and stored at −80°C. Of the specimens used for spatial transcriptomics, tissue samples were obtained from hip (three with RA and three with OA), knee (three with RA and five with OA), and metacarpophalangeal (one with RA); six RA and six OA patients were female, one RA and two OA patients were male (Table S1 for additional information). The mean age was 60.4 ± 5.8 years for the RA patients and 71.6 ± 7.7 years for the OA patients.

### Quality control and sectioning

Frozen tissues were cryosectioned using a Thermo Scientific CryoStar NX70 at −20°C. Slices of 5-μm thickness were transferred to a glass slide. Slides were stored at −80°C before proceeding with staining. The quality and grading of synovitis of the sections were evaluated by histologic analysis (H&E staining). The synovium was identified by the presence of a lining layer and characteristic histologic features of synovium (2).

### GeoMx Digital Spatial Profiler

Sections were thawed at room temperature for 20 min and submerged in 10% NBF (neutral formalin buffer) overnight at room temperature. After washing steps, the sections were incubated with 1X Tris-EDTA (pH 9) at 85 °C for 10min for target retrieval. Sections were washed followed by incubation of proteinase K (1ug/ml) for 15min. A photocleavable RNA probe-set (Human NGS Whole Transcriptome Atlas RNA, NanoString) was applied to the sections and inserted into a hybridization oven at 37 ^°^C overnight. Samples were washed in 50% deionized formamide in sodium citrate buffer (SCC) (ThermoFisher, no. AM9763), then blocked with Buffer W (NanoString) for 30 min. The sections were then incubated for 1 hour with Vimentin (Phycoerythrin; Santa Cruz Biotechnology, no. SC-373717) and CD68 (Alexa 647; Santa Cruz Biotechnology, no. SC-51191), and a nuclear counterstain (Invitrogen, no. S7575).

Slides were washed in SCC buffer and then loaded onto the GeoMx ® Digital Spatial Profiler, and sections were imaged using the ×20 objective. Probes were collected from regions of interest (ROIs) within the synovial lining and sublining through two cell type areas of illumination (AOIs): fibroblast (CD68-Vimentin+), macrophages (CD68+, non-overlapping with “fibroblast” AOIs).

### GeoMx DSP analysis

Sequencing was performed on a NovaSeq 6000 and compiled into FASTQ files. Files were processed to digital count conversion files for each ROI and AOI using the NanoString GeoMX NGS pipeline. Analysis includes ROIs and AOIs that passed manufacturer-recommended quality control thresholds. Segments with <10% of genes detected or <50% sequencing saturation were excluded. Genes detected in <10% of AOIs were removed, except for a list of pathogenic genes, which were considered for analysis regardless of detection level (Table S2). A total of 10,443 genes were retained for downstream analyses. Counts were normalized using quartile three normalization for each AOI. Z scores for selected genes for individual AOIs were used to visualize the expression values.

### Differentially expressed genes and pathway analysis

Samples were separated based on disease (OA or RA), region (lining or sublining), and cell types (fibroblasts or macrophages) to yield unique disease, region, and cell type groups.

Differential expression analysis was performed using linear mixed effects models (LMMs) in RA versus OA (∼ Disease + (1 | TissueSample)) and lining versus sublining (∼ Region + (1 + Region | TissueSample)). Gene-targeting probes with p-value<0.05 and |log2 fold change|>0.5 were defined as differentially expressed genes (DEGs).

For each cell type, all Entrez Gene IDs corresponding to differentially expressed probes (up- regulated and down-regulated) from RA versus OA and lining vs sublining comparisons were submitted to the Reactome Pathway Database using ReactomePA (version 1.50.0) (14) based on Reactome.db version 1.89.0 (15), and clusterProfiler R packages (16). Over-representation analysis was conducted using the hypergeometric distribution, and p-values were adjusted for multiple testing using Benjamini-Hochberg correction. Pathways with a false discovery rate (FDR) <0.02 were considered as being significantly enriched.

## Results

### Experimental overview, cell-type and region-specific gene expression signatures in RA synovium

Fresh frozen synovial tissues from 15 patients (7 RA and 8 OA) were analyzed using the NanoString GeoMx Digital Spatial Profiler (DSP) to investigate spatial and cell-type transcriptional differences in RA and OA (Figure 1A; See Methods). A total of 188 regions of interest (ROIs) were selected from synovial tissue sections, focusing on lining (3-4 ROIs/sample) and sublining (3-4 ROIs/sample) regions in two main cell-type areas AOIs: macrophages (CD68+ Vimentin-) and fibroblasts (CD68- Vimentin+).

**Figure 1.**
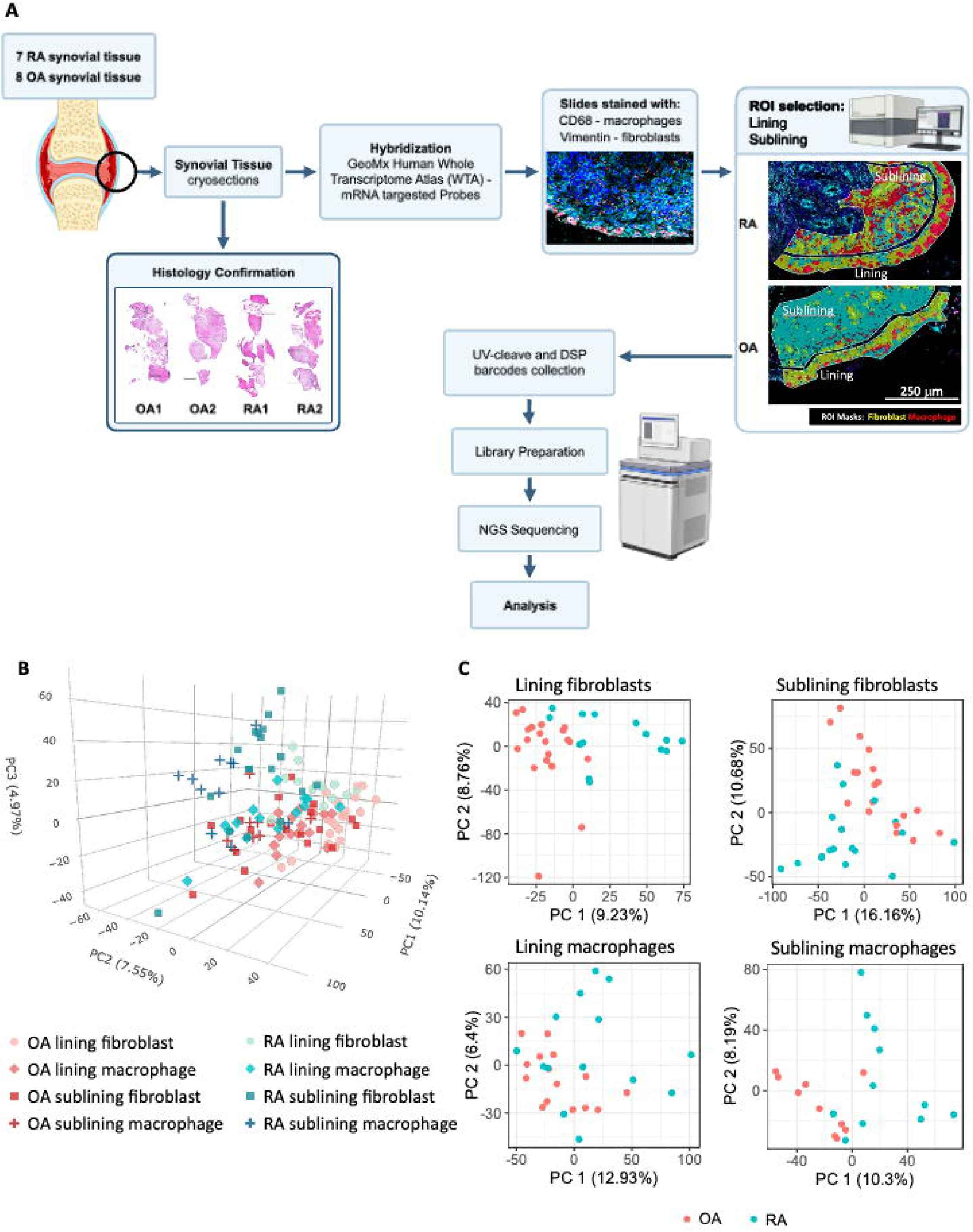
Experimental overview and Principal Component Analysis. **A)** Schematic flowchart of the spatial transcriptomics using the NanoString GeoMx Digital Spatial Profiler. Fresh-frozen synovial tissues from 7 RA and 8 OA patients cut into 5uM sections, followed by H&E histological confirmation, and staining with CD68 in red (macrophages) and vimentin in light blue (fibroblasts). Regions of interest (ROIs) were selected from both the lining and sublining regions based on marker expression. ROIs underwent UV-cleavage to release barcoded oligonucleotides, which were collected and processed for library preparation and next-generation sequencing (NGS). **B)** Principal Component Analysis of normalized gene expression across all segments. Samples were separated by disease (RA vs OA), anatomical region (lining vs sublining), and cell type (fibroblast vs macrophage). **C)** Principal Component Analysis (PCA) of lining fibroblasts (top left) and sublining fibroblasts (top right) regions, visualized by disease status (RA in blue, OA in red). PCA of lining macrophages (bottom left) and sublining macrophages (bottom right) regions are also shown.

The Whole Transcriptome Atlas mRNA probe set was used for hybridization, which provides an unbiased protein-coding gene set. ROI were selected and UV-cleavage was used to release barcoded oligonucleotides, followed by collection, library preparation, and next-generation sequencing (NGS) for digital transcript quantification. After applying QC criteria in both segments and probes, 121 high-quality segments and 10,444 features (including one negative control probe) were retained for downstream analysis.

Principal Component Analysis (PCA) of normalized overall expression data was then performed and showed segregation of segments based on disease state (RA vs OA), anatomical region (lining vs sublining), and cell type, with a focus on fibroblasts and macrophages due to their role in RA and robust staining compared to lymphocytes with this technology (Figure 1B). Even in this complex figure, RA lining fibroblasts and sublining fibroblasts regions could be segregated compared to OA, indicating differences in gene expression related to disease and cell type.

We then performed separate unsupervised clustering analyses by disease and region for each cell type and used PCA to reduce dimensionality and simplify data visualization. This allowed us to see clear differences between cell types and regions. For example, in Figure 1C, PCA revealed prominent separation between RA and OA for lining fibroblasts and sublining fibroblasts, indicating a regional and disease-specific differential gene expression in RA compared to OA. Of interest, lining macrophages exhibited greater similarity between RA and OA than lining fibroblasts, and RA sublining macrophages had overlap with OA (Figure 1C).

### Distinct regional fibroblast transcriptomes in RA compared with OA: Inflammation and matrix remodeling in the lining and adaptive immunity in the sublining

#### Lining fibroblasts: Inflammation and matrix remodeling

To identify gene expression signatures and pathways in synovial fibroblasts, we compared RA and OA regions from the lining and sublining regions separately. Volcano plots revealed distinct sets of DEGs between RA and OA in both regions (Figure 2A). In the lining region, a total of 326 genes were highly expressed in RA fibroblasts. Figure 2B left, shows the top 35 highly expressed DEGs in RA with higher expression of genes involved in extracellular matrix remodeling (e.g., MMP1, MMP3, MMP9, MMP13, TIMP1, CEMIP, COL3A1), inflammatory signaling (e.g., CXCL1, CXCL6, CXCL8, SIK1, CCL18, IL6, S100A9), metabolic stress-related and transcriptional regulators genes (e.g., NR4A1, SLC16A3), and immune modulators such as HLA-DQA2, SPP1, and AIM2. These data indicate an activated, tissue-destructive, and immunomodulatory fibroblast phenotype in the lining region. The DEGs are listed in S3 Table.

**Figure 2.**
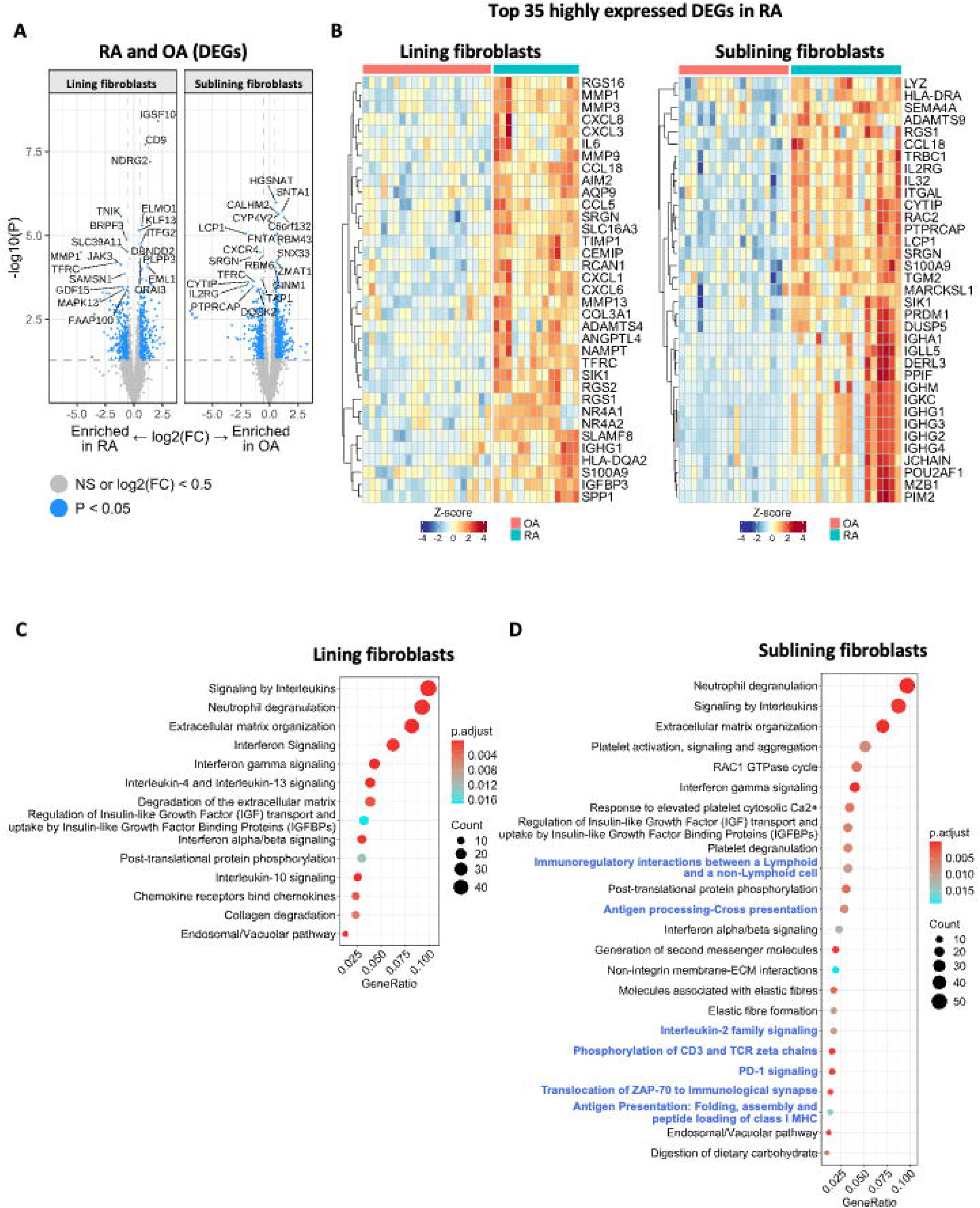
Differentially expressed genes between OA and RA for fibroblasts in the lining and sublining regions. **A)** Volcano plots showing differentially expressed genes between RA and OA fibroblasts in the lining (left) and sublining (right) regions (p-value < 0.05 and |log2(FC)| ≥ 0.5). The x-axis represents the log2 fold change (log2FC) between RA and OA, with values < 0 indicating enrichment in RA and > 0 indicating enrichment in OA. The y-axis represents the - log10 of the p-value, indicating statistical significance. **B)** Heatmaps show the Z score of gene expression level of the top 35 highly expressed DEGs by log2(FC) < 0 in RA fibroblasts from the lining (left) and sublining (right) regions (p-value < 0.05). **C-D)** Dot plots showing Reactome pathway enrichment analysis for DEGs between RA and OA in lining **C)** and sublining **D)** fibroblasts (blue font represents adaptive immune pathways) (FDR<0.02).

Furthermore, the Reactome pathway enrichment analysis of RA versus OA DEGs showed 14 significantly enriched pathways in lining fibroblasts, with strong signals related to inflammation pathways (e.g., interleukins signaling, chemokine receptor bind chemokines, interferon signaling, post-translational protein phosphorylation), and matrix regulation pathways (e.g., extracellular matrix organization, degradation of the extracellular matrix, collagen degradation) (Figure 2C). The clustered DEGs from inflammation or matrix regulation pathways revealed that most were highly expressed in RA (S1 Figure). Among these included MMP1, MMP3, CXCL1, CXCL6, IL6, and TIMP1.

#### Sublining Fibroblasts: Inflammation and adaptive immunity

In the sublining region, 277 DEGs were highly expressed in RA fibroblasts, with the top 35 included genes associated with immune activation (e.g., IL32, CCL18), regulators of cell survival and stress response (e.g., PIM2, TGM2, RAC2), MHC class II–related molecules (HLA-DRA), and immunoglobulin-related genes (Figure 2B right). The DEGs with fold-change values and P-values are listed in S3 Table.

The pathway enrichment analysis in sublining fibroblasts from RA versus OA DEGs showed enrichment for 24 pathways, some of which were similar to lining fibroblasts, such as some inflammatory and matrix regulation pathways (e.g., extracellular matrix organization, interleukin signaling, interferon signaling) (Figure 2D). Unexpectedly, pathways related to the regulation of adaptive immunity, antigen presentation, and lymphocyte activation were also identified in the sublining fibroblast transcriptional profile (e.g., immune regulatory interactions between a lymphoid and a non-lymphoid cell, antigen processing-cross presentation, IL2 signaling, PD1 signaling, and antigen presentation). Clustering of DEGs in the adaptive immune cell signaling– related pathways revealed that most were highly expressed in RA, these included HLA genes and IL2RG, CD8A, B2M, ITGAL, and PTPRC (S2 Figure).

In summary, our findings demonstrate that RA fibroblasts display region-specific transcriptional programs, with lining fibroblasts exhibiting a more matrix-destructive and pro-inflammatory phenotype, while sublining fibroblasts are more involved in communication between the immune system and antigen processing.

### Distinct regional macrophage transcriptomes in RA compared with OA

#### Lining macrophages: Inflammation and Metabolic Activation

We then compared RA and OA macrophages within the lining and sublining regions. Volcano plots showed different sets of DEGs in both regions (Figure 3A). A total of 315 highly expressed DEGs in RA lining macrophages were identified. Of interest, some of these genes overlapped with those found in RA lining fibroblasts from the same region, including in extracellular matrix remodeling (e.g., MMP1, MMP3, MMP13, TIMP1), and inflammatory signaling (e.g., CXCL1, CXCL3, CCL18), suggesting a shared inflammatory and tissue-destructive phenotype across cell types (Figure 3B, left). However, they also showed a unique expression profile in RA lining, including immunoglobulin-related genes (e.g., IGHG2, IGHG3), extracellular matrix remodeling (e.g., MMP19, COL1A1, COL5A3), and innate immune regulators (e.g., LBP, CHI3L1), which were not prominently expressed in RA lining fibroblasts. Thus, lining macrophages transcriptomes suggest that they participate in tissue remodeling and increased responsiveness to pathogens or damage within the RA synovial environment.

**Figure 3.**
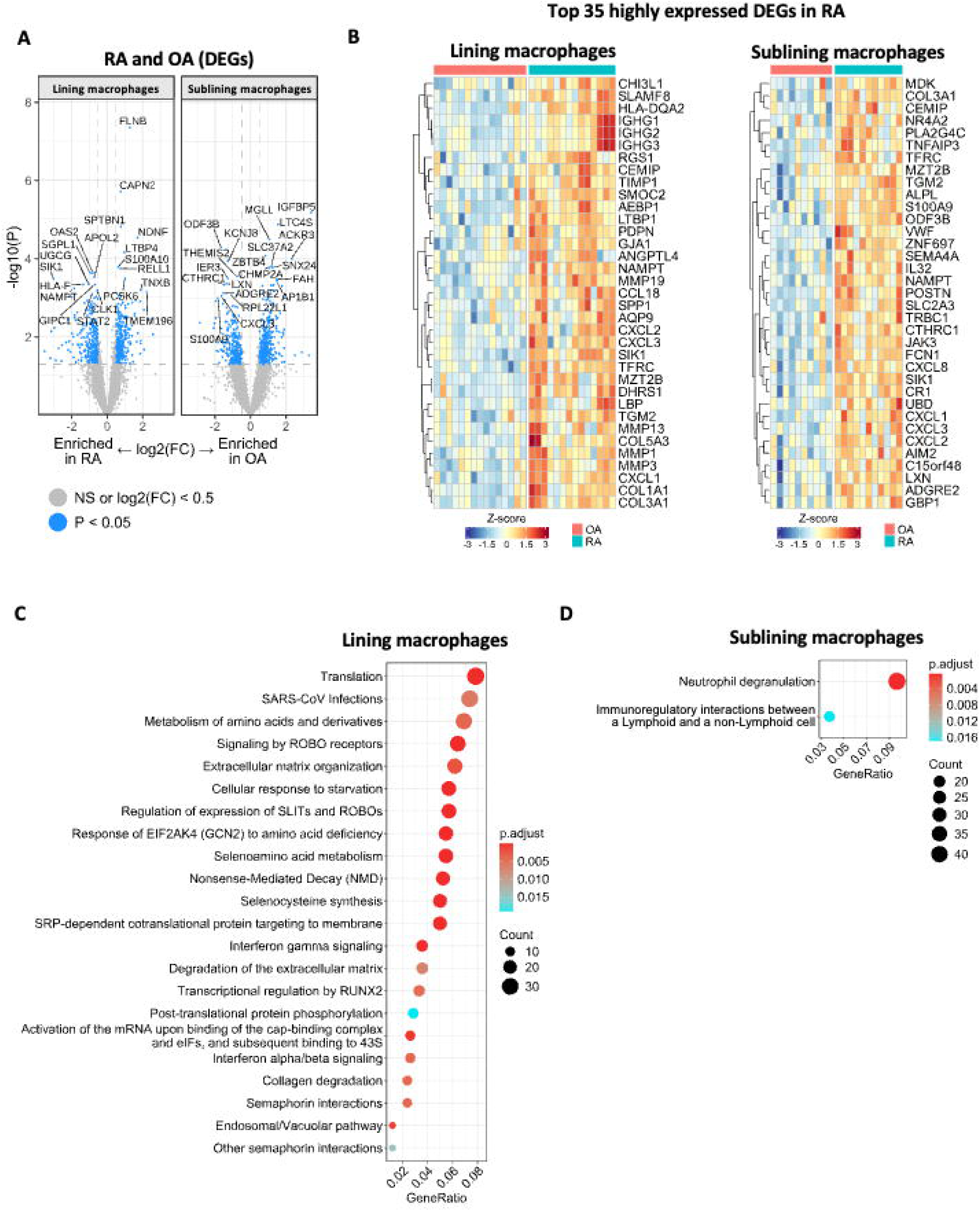
Differentially expressed genes between OA and RA for macrophages in the lining and sublining regions. **A)** Volcano plots showing differentially expressed genes between RA and OA macrophages in the lining (left) and sublining (right) regions (p-value < 0.05 and |log2(FC)| >= 0.5). The x-axis represents the log2 fold change (log2FC) between RA and OA, with values < 0 indicating enrichment in RA and > 0 indicating enrichment in OA. The y-axis represents the - log10 of the p-value, indicating statistical significance. **B)** Heatmaps show the Z score of gene expression of the top 35 highly expressed DEGs by log2(FC) < 0 in RA macrophages from the lining (left) and sublining (right) regions (p-value < 0.05). **C-D)** Dot plots of Reactome pathway over-representation analysis for lining **(C)** and sublining **(D)** RA macrophages (FDR<0.02). A full list of enriched pathways for lining **(C)** is provided in S2 Figure.

Pathway enrichment analysis of RA versus OA DEGs revealed 47 significantly enriched pathways (Figure 3C and S3), mostly related to translation and ribosomal processes (e.g., cap-dependent translation initiation, rRNA processing), amino acid metabolism, interferon signaling, and extracellular matrix organization. Therefore, lining macrophages exhibit genes involved with increased metabolic and biosynthetic activity, supporting local inflammation and matrix remodeling.

#### Sublining macrophages: Lack of differential gene expression in RA synovium

Sublining macrophages had 268 DEGs highly expressed in RA. The top 35 highly expressed genes were involved in inflammatory activation, such as SIK1, CXCL2, and CXCL1, in neutrophil recruitment and innate immune activation (e.g., SEMA4A and S100A9), and some related to vascular interaction and complement immune regulation (e.g., VWF, TFRC). The DEGs are listed in S3 Table.

However, pathway enrichment analysis of RA versus OA DEGs revealed different biological profiles between lining and sublining macrophages. While lining macrophages were significantly different in RA in numerous pathways (Fig 3C), sublining macrophages showed enrichment of only 2 pathways (neutrophil degranulation and immunoregulatory interactions between lymphoid and non-lymphoid cells) (Figure 3D). The paucity of pathways in sublining macrophages suggests that they do not drive RA pathology.

### Potential roles of lining fibroblasts and lining macrophages in RA pathogenesis

To visualize differences in transcriptome profiles between cell types and regions, we developed a Venn diagram to compare DEGs between RA and OA (Figure 4A). A total of 700 DEGs were found in lining fibroblasts (334 unique to this group), 848 in sublining fibroblasts (496 unique), 627 in lining macrophages (369 unique), and 673 in sublining macrophages (470 unique). The overlap between DEGs across regions was small, underlining spatial heterogeneity. 38 DEGs were shared across all cell types and regions (S4 Figure), including SIK1, CXCL2, CXCL3, and JAK3. Those genes were highly expressed in RA, consistent with their known roles in inflammatory signaling, and others, such as TMEM196, NDNF, THBS4, and COLEC12, were higher in OA. However, no enrichment Reactome pathways were found from these 38 DEGs, possibly due to the low number of genes in the list.

**Figure 4.**
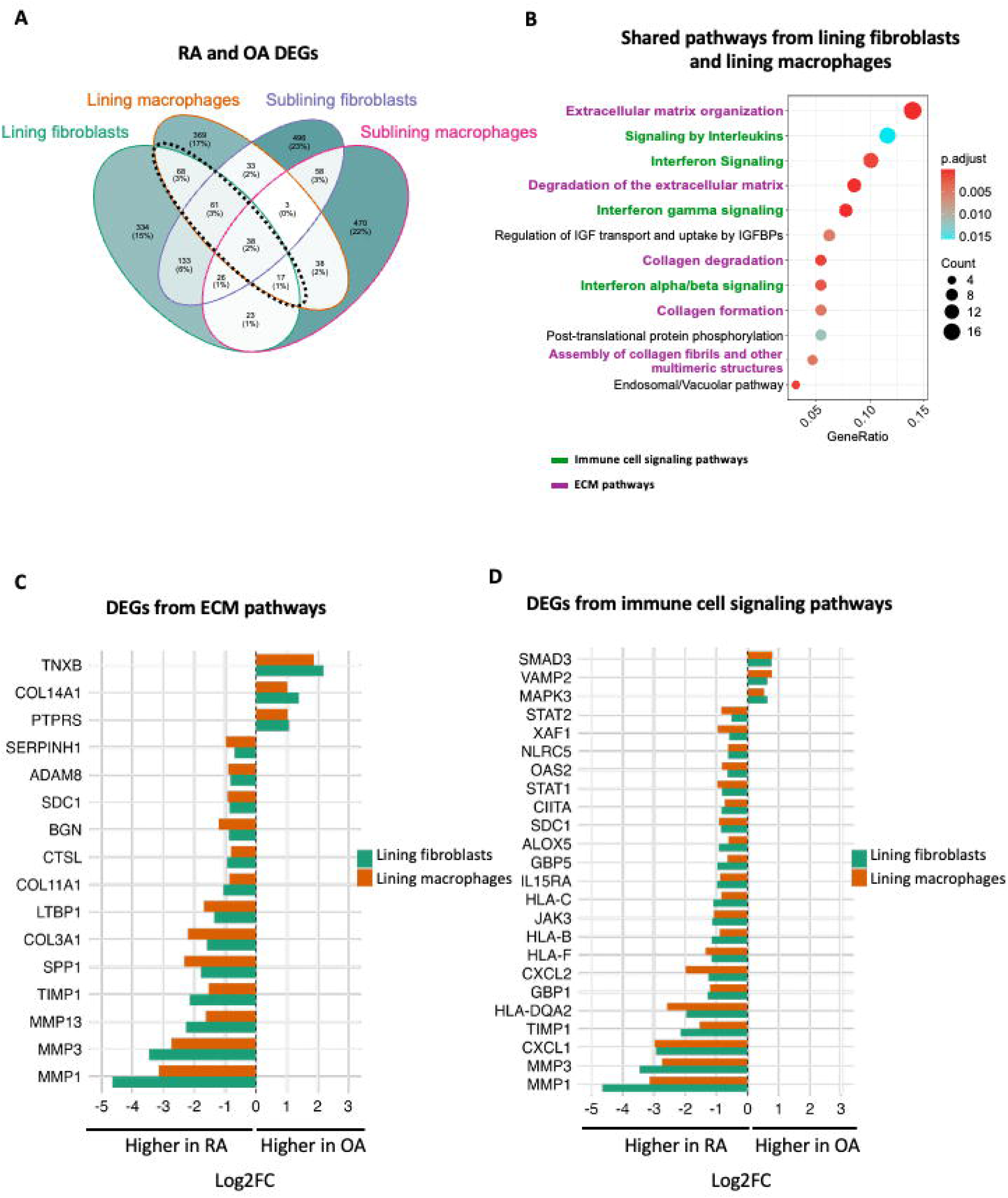
Shared differentially expressed genes (DEGs) and enriched pathways in RA vs OA for synovial lining fibroblasts and lining macrophage. **A)** Venn diagram showing the number of DEGs in lining fibroblasts (green) and lining macrophages (orange) from RA vs OA. The dotted bold lines show the overlap genes between the two cell types, revealing 184 shared DEGs. **B)** Dot plot of Reactome pathway enrichment analysis for the 184 overlapping DEGs in lining fibroblasts and lining macrophages (FDR <0.02). Pathways are color-coded by relation to ECM pathways (purple) or immune cell signaling (green). **C-D)** Bar-plots show the Log2(FC) (RA vs OA) of DEGs in lining fibroblasts and lining macrophages found in ECM pathways **(C)** or immune cell signaling **(D)**. The negative log2(FC) shows highly expressed genes in RA, and the positive log2(FC) indicates highly expressed genes in OA.

Given that fibroblasts and macrophages localize to the synovial lining, a critical region that drives RA pathogenesis (6), we also investigated their common gene expression patterns. In both cells, we found 184 overlapping DEGs in RA versus OA, with 92 genes highly expressed in RA (Figure 4A and S3 Table). Among these genes involved in ECM remodeling (MMP1, MMP3, MMP13), immune regulation (CEMIP, SPP1, SIK1), and chemokine signaling (CXCL1, CCL18, CXCL3).

The pathway analysis of the shared DEGs showed 12 commonly enriched pathways, including ECM organization and degradation, interleukin and interferon signaling, and collagen formation (Figure 4B). Notably, the DEGs clustered in ECM and immune signaling pathways showed higher expression in RA than in OA (Figure 4C and D), supporting the concept of an aggressive phenotype shared between both cells in RA lining.

### Topographical transcriptome profiling of macrophages and fibroblasts within RA synovium

To understand the region-specific functional specialization within in RA synovium, we also performed differential gene expression comparing lining and sublining regions in fibroblasts and macrophages from RA. In Figure 5A, the volcano plot showed different sets of DEGs enriched in sublining fibroblasts or lining fibroblasts in RA. In total, RA lining fibroblasts showed 553 highly expressed DEGs, including PRG4, SKP1, MMP3, MMP1, and CLU in the top 35 highest ones (Figure 5A, right). Interestingly, these same genes were also among the top highly expressed in RA lining macrophages (Figure 5C). These findings suggest coordinated regulation of joint lubrication and tissue remodeling across cell types in the synovial lining in RA. In addition, RA fibroblasts showed 32 pathways enriched in RHO/RAC1 GTPase cycle, disease metabolism, disease of glycosylation, degradation of extracellular matrix, complement cascade, and TGFß receptor signaling (Figure 5B). These reinforce the idea that lining fibroblasts in RA are highly aggressive.

**Figure 5.**
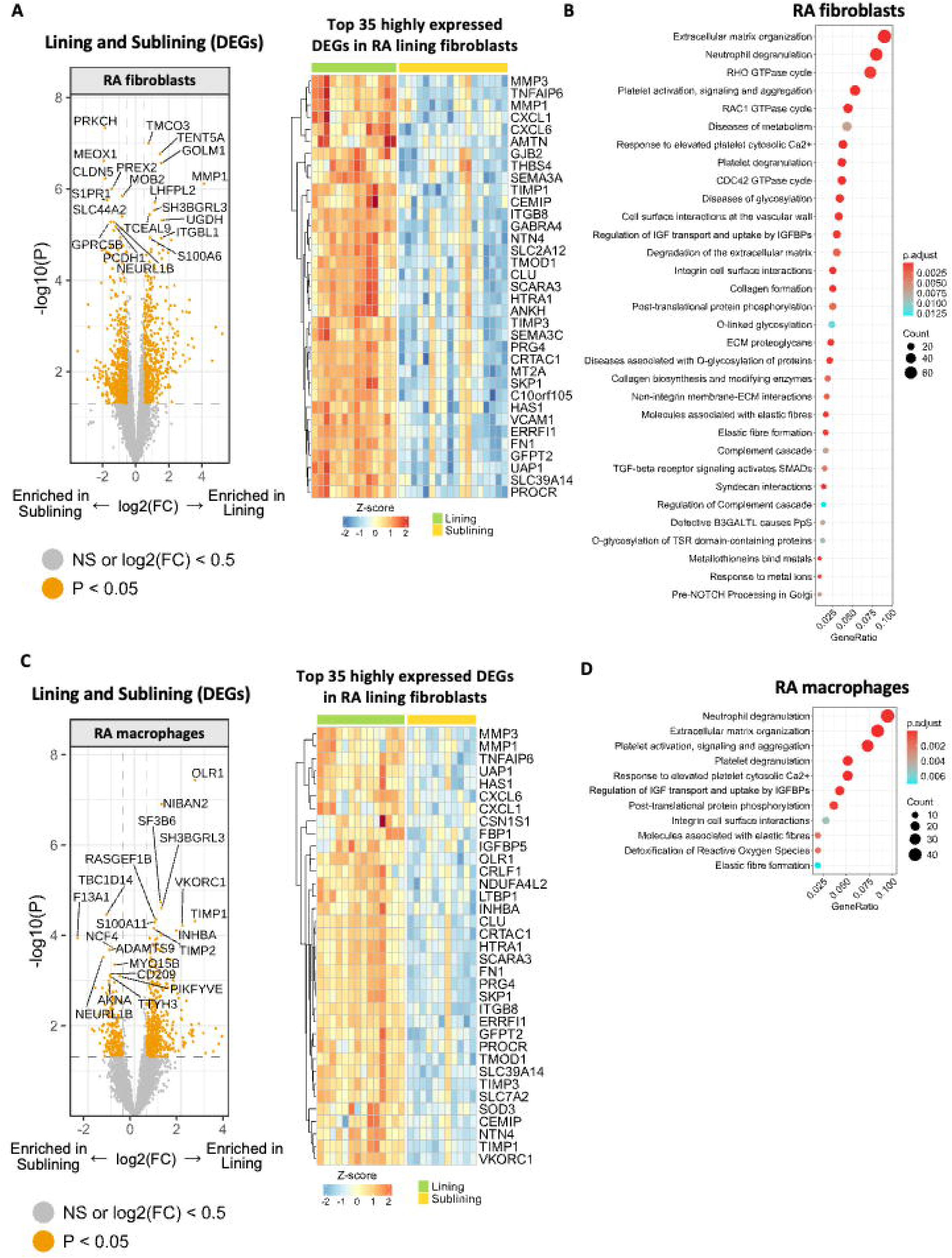
Lining vs sublining gene expression and pathways in RA fibroblasts and macrophages. **A)** Volcano plot showing differentially expressed genes (DEGs) between RA lining and sublining fibroblasts (left). The x-axis represents the log2 fold change (log2FC) between lining and sublining, with values < 0 indicating enrichment in sublining and > 0 indicating enrichment in lining. The y-axis represents the -log10 of the p-value, indicating statistical significance. On the right, the heatmap shows the Z score of expression levels of the top 35 highly expressed DEGs by log2(FC) > 0 in RA lining fibroblasts (p-value < 0.05). **B)** Dot plot showing Reactome pathway enrichment for DEGs in RA fibroblasts (lining vs sublining) (FDR <0.02). **C)** Volcano plot of DEGs between RA lining and sublining macrophages (left), and heatmap of Z score of expression levels of top 35 highly expressed genes by log2(FC) > 0 in RA lining macrophages (right) (p-value < 0.05). **D)** Pathway enrichment analysis for DEGs in RA macrophages (lining vs sublining) (FDR<0.02).

Pathway enrichment analysis of RA macrophage DEGs for lining compared with sublining revealed 11 pathways in RA macrophages, many of which overlapped with RA fibroblasts analysis, including extracellular matrix, post-translational protein phosphorylation, and integrin cell surface interactions (Figure 5D).

## Discussion

Rheumatoid synovium is a site of robust adaptive and innate immune activation (1, 2, 6). There have been dramatic advances in our understanding of RA pathogenesis, but precise synovial mechanisms of disease remain incomplete. The architecture of the joint has been defined based on histopathologic studies and refined with early immunostaining and later *in situ* hybridization and laser capture microscopy (LCM) to map key mediators and cell types to tissue (10–12).

Evaluation of cells isolated from dispersed synovium also revealed critical information on aberrant function, such as the unique aggressive behavior of lining fibroblasts (17–19). The more recent advent of single-cell and unbiased omics technologies has advanced our understanding of isolated synovial cells, but does not offer information on the topographical relationships between cells, and is confounded by artifacts due to the tissue disaggregation.

In the present study, we build on this knowledge with the first broad-based unbiased spatial transcriptomics analysis of RA and OA synovium that helps define cell types and topography that contribute to pathogenesis. Previous spatial studies, including our own, had smaller sample sizes or lacked OA control synovium for comparison to identify DEGs (20). We used GeoMx technology (13, 21), which does not provide single-cell precision but evaluates the entire transcriptome (rather than being probe-based) in combination with immunostaining to define cell lineage. We focused this study on macrophages and fibroblasts because of their important contribution to RA mechanisms and the feasibility related to staining intensity for these cells using this technology. Our results provide a comprehensive regional map of RA synovial macrophage and fibroblast transcriptomes.

The synovial lining has long been known to be highly activated with high expression of MMPs, cytokines like IL-6, prostanoids and other mediators (3, 6). The methodology used to localize these pathogenic mediators mainly relied on techniques like immunostaining and in situ hybridization (22). Recent studies using LCM with RNAseq comparing RA to OA confirmed those results, although the number of DEGs and pathways was much lower than our spatial transcriptomics data (12). It has been difficult to dissect the cellular origin of these products because of the technologies used.

The spatial data now allow us to define gene expression profiles of the two cell types in the lining region, namely macrophages and fibroblasts. Our regional analysis from RA synovium extends previous findings, showing that lining fibroblasts are highly activated in pathways related to immune cell signaling and ECM organization compared to sublining fibroblasts as well as the OA lining (4, 10, 17, 23–25). For example, lining fibroblasts highly express multiple MMPs, integrins, and ADAMs, supporting enrichment for the ECM and collagen degradation pathways. Additionally, they showed strong enrichment for pathways like O-linked glycosylation, RHO GTPase cycle, and complement cascade regulation, with higher expression of genes like GALNT5, FN1, and CD55. In the disease-specific comparison analysis, ECM organization and degradation pathways are strongly enriched in RA lining fibroblasts, including several MMPs, CEMIP, and ICAM1. They also showed marked upregulation of inflammatory mediators such as IL-6, CCL5, CXCL1, CXCL8, and CXCL6, consistent with interleukins signaling and chemokine receptor bind chemokines pathways. These findings also demonstrate that immune activation and tissue destruction of RA lining fibroblasts are consistent with studies showing their aggressive phenotype *in vitro* (18, 19, 23, 24, 26). This provides support for the utility of studies using cultured lining fibroblasts as a model for tissue fibroblasts.

While the data lining fibroblasts expand our understanding of their activation state and transcriptional profile, the most surprising finding is related to RA sublining fibroblasts compared to OA, which implicates their role in adaptive immunity based on DEGs in that region. Genes like HLA family genes, B2M, CTSS, and TAP1/2 were highly expressed by these RA cells and explain sublining fibroblast enrichment of pathways like antigen processing-cross presentation, immunoregulatory interactions between a lymphoid and a non-lymphoid cell, and PD-1 signaling. This observation is consistent with previous reports that defined a potential role of synovial fibroblasts, indicating that IFNγ-treated cultured lining fibroblasts could present antigens to T cells (5, 27).

In addition to adaptive immunity functions, RA sublining fibroblasts also showed enrichment for interleukins signaling, ECM organization, and RAC1 GTPase cycle pathways compared to OA, driven by DEGs like IL32, JAK3, ICAM1/3, ADAMTS9, and TAGAP. Previous studies have defined a unique phenotype of sublining synovial fibroblasts based on presence of proteins like CD90 and FAPα (18). Our findings are consistent with immunostaining and studies using isolated sublining fibroblasts, suggesting that they possess pro-inflammatory functions (28–30). This phenotypic continuum appears to be driven by the microenvironment, such as local expression of NOTCH3 by endothelial cells (28, 30, 31). Thus, it is likely that the sublining fibroblast cells are plastic rather than terminally differentiated and, like synovial macrophages (32), can alter their inflammatory and adaptive immunity functions as they migrate from perivascular regions to the lining.

The macrophage spatial transcriptomics data are also illuminating. Lining macrophages express a large number of DEGs consistent with metabolic and immune activation. Among these DEGs, a few immunoglobulin-related genes were also detected, consistent with previous studies (33, 34). DEGs like MRLP14, HLA family genes, and SLC7A5 (LAT1) were upregulated in RA lining macrophages compared to OA. The overall pattern shows that these cells have an increased metabolic and biosynthetic activity. RA lining macrophages also showed a strong enrichment for ECM organization pathways compared to OA, including DEGs such as MMPs, TIMPs, and ADAMs. However, the most striking observation for macrophages was the lack of DEGs and pathways comparing RA and OA sublining macrophages. This unexpected finding suggests that sublining macrophages are not a distinguishing feature or pathogenic driver of rheumatoid synovitis. This rather quiescent transcriptome contrasts with lining macrophages or fibroblasts in either region.

Our study does have some limitations. While we had sufficient material to identify critical pathways, the number of DEGs could be underestimated due to the sample size. This, along with the fact that our samples are largely de-identified, also limits our ability to make clinical correlations. We also infer functional properties based on the transcriptome without *in vitro* biological validation. In addition, multiple DEGs expressed by fibroblasts and macrophages in our analysis were shown in pathways such as neutrophil degranulation, suggesting the pathway analysis should be interpreted with some caution. The process of immunostaining and spatial transcriptomics makes direct studies of the identified cells impossible. However, future studies looking at the function of disaggregated cells might address this need. Also, while single-cell resolution would be important for evaluating cellular niches, GeoMx does not achieve true single-cell resolution. Future studies using orthogonal single-cell technologies will be needed to validate these findings. Nevertheless, the immunofluorescence-based approach of GeoMx offers the advantage of defining cell lineage based on protein rather than mRNA.

In conclusion, we showed specific transcriptional profiles in RA synovium based on cell lineage and location. These data provide clues on the relative contributions of each cell type to the pathogenesis of disease. Surprising findings included the involvement of sublining fibroblasts in adaptive immunity and the lack of significant pathway enrichment for sublining macrophages.

Potential therapeutic targets also emerge based on targeting specific regions, cell phenotypes, and DEGs. Future studies integrating additional technologies, such as ATACseq or CITEseq, as well as ongoing single spatial cell studies using probe-based approaches, will hopefully continue to expand the synovial gene expression map.

## Acknowledgements

We wish to acknowledge the patients who provided synovial tissues, the team of the Division of Rheumatology, Autoimmunity and Inflammation, and Eli Lilly and Company for funding the study and contributing to its design, sample collection, and data interpretation.

## Data availability

The data will be deposited in public databases and will be made available upon publication.

## References

1. Firestein GS. Evolving concepts of rheumatoid arthritis. Nature2003.

2. Veale D, and GS Firestein. Synovium, In: Firestein and Kelley’s Textbook of Rheumatology, GS Firestein, et al. eds., Elsevier, Philadelphia, 11th edition, 2020.

3. Bartok B, Firestein GS. Fibroblast-like synoviocytes: key eUector cells in rheumatoid arthritis. Immunol Rev. 2010;233(1):233–55.

4. Smith MH, Gao VR, Periyakoil PK, et al. Drivers of heterogeneity in synovial fibroblasts in rheumatoid arthritis. Nat Immunol. 2023;24(7):1200–10.

5. Tran CN, Lundy SK, Fox DA. Synovial biology and T cells in rheumatoid arthritis. Pathophysiology. 2005;12(3).

6. Firestein GS, McInnes IB. Immunopathogenesis of Rheumatoid Arthritis. Immunity. 2017;46(2):183–96.

7. Tu J, Hong W, Zhang P, et al. Ontology and Function of Fibroblast-Like and Macrophage-Like Synoviocytes: How Do They Talk to Each Other and Can They Be Targeted for Rheumatoid Arthritis Therapy? Front Immunol. 2018;9:1467

8. Culemann S, Grüneboom A, Nicolás-Ávila JÁ, et al. Locally renewing resident synovial macrophages provide a protective barrier for the joint. Nature. 2019;572(7771).

9. Boyle DL, Prideaux EB, Hillman J, et al. Improving Transcriptome Fidelity Following Synovial Tissue Disaggregation. Frontiers in Medicine. 2022;9.

10. Zhang F, Wei K, Slowikowski K, Fonseka CY, Rao DA, Kelly S, et al. Defining inflammatory cell states in rheumatoid arthritis joint synovial tissues by integrating single-cell transcriptomics and mass cytometry. Nature Immunology. 2019;20(7).

11. Zhang F, Jonsson AH, Nathan A, et al. Deconstruction of rheumatoid arthritis synovium defines inflammatory subtypes. Nature. 2023;623(7987).

12. Van Espen B, Prideaux EB, Wilson AR, et al. Laser Capture Microscopy RNA Sequencing for Topological Mapping of Synovial Pathology During Rheumatoid Arthritis. Arthritis and Rheumatology. 2024;76(8):1243–51.

13. Dong Y, Saglietti C, Bayard Q, et al. Transcriptome analysis of archived tumors by Visium, GeoMx DSP, and Chromium reveals patient heterogeneity. Nat Commun. 2025;16(1):4400.

14. Yu G, He QY. ReactomePA: An R/Bioconductor package for reactome pathway analysis and visualization. Molecular BioSystems. 2016;12(2).

15. Milacic M, Beavers D, Conley P, et al. The Reactome Pathway Knowledgebase 2024. Nucleic Acids Research. 2024;52(D1).

16. Yu G, Wang LG, Han Y, et al. ClusterProfiler: An R package for comparing biological themes among gene clusters. OMICS A Journal of Integrative Biology. 2012;16(5).

17. Mizoguchi F, Slowikowski K, Wei K, et al. Functionally distinct disease-associated fibroblast subsets in rheumatoid arthritis. Nature Communications. 2018;9(1).

18. Qian H, Deng C, Chen S, et al. Targeting pathogenic fibroblast-like synoviocyte subsets in rheumatoid arthritis. Arthritis Research and Therapy: BioMed Central Ltd; 2024.

19. Machado CRL, Choi E, Perumal NB, et al. Regulation of fibroblast-like synoviocyte function by cadherin 6 in rheumatoid arthritis. Arthritis Research and Therapy. 2025;27(1).

20. Vickovic S, Schapiro D, Carlberg K, et al. Three-dimensional spatial transcriptomics uncovers cell type localizations in the human rheumatoid arthritis synovium. Communications Biology. 2022;5(1).

21. Cho C, Haddadi NS, Kidacki M, et al. Spatial Transcriptomics in Inflammatory Skin Diseases Using GeoMx Digital Spatial Profiling: A Practical Guide for Applications in Dermatology. JID Innov. 2025;5(1):100317.

22. Firestein GS, Paine MM, Boyle DL. Mechanisms of Methotrexate Action in Rheumatoid-Arthritis - Selective Decrease in Synovial Collagenase Gene-Expression. Arthritis and Rheumatism. 1994;Vol 37(Iss 2).

23. Ainsworth RI, Hammaker D, Nygaard G, et al. Systems-biology analysis of rheumatoid arthritis fibroblast-like synoviocytes implicates cell line-specific transcription factor function. Nature Communications. 2022;13(1).

24. Choi E, Machado CR, Okano T, et al. Joint-specific rheumatoid arthritis fibroblast-like synoviocyte regulation identified by integration of chromatin access and transcriptional activity. JCI Insight. 2024;9(12).

25. Rauber S, Mohammadian H, Schmidkonz C, et al. CD200+ fibroblasts form a pro-resolving mesenchymal network in arthritis. Nature Immunology. 2024;25(4).

26. Knab K, Chambers D, Krönke G. Synovial Macrophage and Fibroblast Heterogeneity in Joint Homeostasis and Inflammation. Frontiers in Medicine. 2022;9:862161.

27. Tran CN, Davis MJ, Tesmer LA, et al. Presentation of arthritogenic peptide to antigen-specific T cells by fibroblast-like synoviocytes. Arthritis Rheum. 2007;56(5):1497–506.

28. Croft AP, Campos J, Jansen K, et al. Distinct fibroblast subsets drive inflammation and damage in arthritis. Nature. 2019;570(7760):246–51.

29. Mizoguchi F, Slowikowski K, Wei K, et al. Functionally distinct disease-associated fibroblast subsets in rheumatoid arthritis. Nat Commun. 2018;9(1):789.

30. Wei K, Korsunsky I, Marshall JL, Gao A, Watts GFM, Major T, et al. Notch signalling drives synovial fibroblast identity and arthritis pathology. Nature. 2020;582(7811):259–64.

31. Marsh LJ, Kemble S, Reis Nisa P, et al. Fibroblast pathology in inflammatory joint disease. Immunol Rev. 2021;302(1):163–83.

32. Hogg N, Palmer DG, Revell PA. Mononuclear phagocytes of normal and rheumatoid synovial membrane identified by monoclonal antibodies. Immunology. 1985;56(4).

33. Gong X, Yan H, Ma J, Zhu Z, et al. Macrophage-Derived Immunoglobulin M inhibits inflammatory responses via modulating Endoplasmic reticulum stress. Cells. 2021;10(11).

34. Fuchs T., Hahn M., Ries L., et al. Expression of combinatorial immunoglobulins in macrophages in the tumor microenvironment. PLoS One. 2018;13:e0204108.

